# Regulatory interactions between daptomycin and bacitracin responsive pathways coordinate the cell envelope antibiotic resistance response of *Enterococcus faecalis*

**DOI:** 10.1101/2022.11.16.516778

**Authors:** Sali M. Morris, Laura Wiens, Olivia Rose, Georg Fritz, Tim Rogers, Susanne Gebhard

## Abstract

Enterococcal infections frequently show high levels of antibiotic resistance, including to cell envelope-acting antibiotics like daptomycin (DAP). While we have a good understanding of the resistance mechanisms, less is known about the control of such resistance genes in enterococci. Previous work unveiled a bacitracin resistance network, comprised of the sensory ABC transporter SapAB, the two-component system (TCS) SapRS and the resistance ABC transporter RapAB. Interestingly, components of this system have recently been implicated in DAP resistance, a role usually regulated by the TCS LiaFSR. To better understand the regulation of DAP resistance and how this relates to mutations observed in DAP-resistant clinical isolates of enterococci, we here explored the interplay between these two regulatory pathways. Our results show that SapR regulates an additional resistance operon, *dltXABCD*, a known DAP resistance determinant, and show that LiaFSR regulates the expression of *sapRS*. This regulatory structure places SapRS target genes under dual control, where expression is directly controlled by SapRS, which itself is up-regulated through LiaFSR. The network structure described here shows how *E. faecalis* coordinates its response to cell envelope attack and can explain why clinical DAP resistance often emerges via mutations in regulatory components.

## Introduction

The rise of antibiotic resistant bacteria is one of the greatest current threats to public health, resulting in 670,000 infections a year and 33,000 deaths in Europe alone (Organisation for Economic Co-operation and Development, 2020). Of these infections, the “ESKAPE” organisms (Zhen *et al*., 2019) (*Enterococci* spp., *Staphylococcus aureus*, *Klebsiella pneumoniae*, *Acinetobacter baumannii*, *Pseudomonas aeruginosa*, *Enterobacter* spp.) have driven the rising number of nosocomial and antibiotic-resistant infections in the past decade. Of these bacteria, the enterococci are the second most common causative agent of nosocomial infections in the US; including bacteraemia, endocarditis and urinary tract infections (Miller *et al*., 2016; Chiang *et al*., 2017; Said *et al*., 2021). The two species most frequently isolated, *Enterococcus faecalis* and *Enterococcus faecium*, remain a major infection-control challenge, particularly in healthcare settings.

Enterococci became recognised as important nosocomial pathogens due to their high level of intrinsic resistance to several antimicrobials (Arias and Murray, 2012) (e.g. penicillin, ampicillin and cephalosporins) and their capacity to acquire further resistance determinants. One such acquired resistance is to the glycopeptide antibiotic vancomycin, which occurs through plasmid acquisition and was first reported in the 1988 (Leclercq *et al*., 1988; Uttley *et al*., 1989), 30 years after vancomycin was introduced for clinical use (Levine, 2006). Despite the molecular mechanisms of vancomycin resistance in enterococci being well understood today (Courvalin, 2006), infections by vancomycin resistant enterococci (VRE) still result in serious health and economic impacts (Puchter *et al*., 2018) and are an increasing problem worldwide.

One of the last-resort antibiotics used to treat these VRE infections is the lipopeptide antibiotic daptomycin (DAP) (Sauermann *et al*., 2008). Disappointingly, within two years of clinical introduction of the drug in 2003, DAP-resistant enterococcal isolates were reported (Munoz-Price *et al*., 2005), and in contrast to vancomycin resistance, this occurred through a subtle chromosomal change based on the mutations in genes *liaF*, *cls* and *gdpD* (Palmer *et al*., 2011; Arias *et al*., 2011; Tran *et al*., 2013; Reyes *et al*., 2015). LiaF is a transmembrane protein involved in monitoring the integrity of the cell envelope and responding to damage (Tran *et al*., 2016), whereas Cls (cardiolipin synthase) and GdpD (glycerol-phosphodiester phosphodiesterase) are both involved in phospholipid metabolism (Arias *et al*., 2011). Of these, mutation of *liaF* has been proposed to be the first pivotal event towards DAP resistance (Miller *et al*., 2013; Tran *et al*., 2013). However, the exact role of the *liaF* mutations in DAP resistance so far remains unclear.

LiaF is part of a three-component regulatory system, LiaFSR, which is important amongst the Firmicutes for coordinating the cell envelope stress response (CESR) against antimicrobial-induced damage (Kuroda *et al*., 2003; Jordan *et al*., 2006; Martínez *et al*., 2007; Suntharalingam *et al*., 2009). The system is comprised of LiaF and a conventional two-component system (TCS): the sensor kinase LiaS and the response regulator LiaR (Jordan *et al*., 2006; Schrecke *et al*., 2013; Wolf *et al*., 2010). LiaF is an inhibitor of LiaS, maintaining the sensor kinase in an inactive conformation (Jordan *et al*., 2006). Rather than detecting a specific antimicrobial compound, LiaFSR responds to cell envelope damage, although the exact stimulus is unknown (Wolf *et al*., 2012). Upon sensing this damage, LiaF releases LiaS, which is then able to phosphorylate LiaR to induce the expression of the system’s target operon, *liaXYZ* (Khan *et al*., 2019; Miller *et al*., 2013). The *liaXYZ* cluster is thought to be involved in sensing and binding antimicrobials at the cell surface to provide resistance (Khan *et al*., 2019a).

To add to this complexity, the Lia system does not exist in isolation, but is just one of many TCSs involved in monitoring cell envelope integrity, each responding to their own individual stimuli and activating a unique set of target genes. The CroRS system, unique to the enterococci, is the main regulator required for cephalosporin resistance (Comenge *et al*., 2003), whereas the VicKR (YycFG) system is essential across the low-CG Gram positives and is involved in regulating cell division, lipid biosynthesis, biofilm and virulence (Winkler and Hoch, 2008). An additional element of the network is the serine/threonine kinase IreK and the phosphatase IreP, involved in maintaining cell wall integrity by potentially regulating peptidoglycan biosynthesis and metabolism (Kristich *et al*., 2007a; Kristich *et al*., 2011; Iannetta *et al*., 2021).

A further TCS involved in monitoring the cell envelope is EF0926/27, which we have now renamed SapRS (<u>S</u>ensor of <u>A</u>ntimicrobial <u>P</u>eptides), identified in our previous work (Gebhard *et al*., 2014). SapRS is part of a bacitracin (BAC) resistance network comprised of the histidine kinase SapS, the response regulator SapR and the Bce-like ABC transporters: SapAB (EF2752/51) and RapAB (<u>R</u>esistance against <u>A</u>ntimicrobial <u>P</u>eptides) (EF2050/49) (Gebhard *et al*., 2014). The use of the sensory transporter SapAB to control the activity of SapRS implements a ‘flux-sensing’ mechanism to regulate BAC resistance, as was shown for the homologous system in *Bacillus subtilis* (Fritz *et al*., 2015). In brief, upon exposure, BAC binds to its membrane-associated target molecule, undecaprenol-pyrophosphate (UPP), blocking the dephosphorylation and recycling of UPP in the lipid II cycle of cell wall biosynthesis (Storm and Strominger, 1974; Storm, 1974). The sensory ABC transporter SapAB, based on biochemical and structural evidence from the *B. subtilis* system, forms a sensory complex with SapRS (Dintner *et al*., 2014; George and Orlando, 2023). In the absence of BAC, the transporter maintains the histidine kinase, SapS, in an ‘OFF’ state (Koh *et al*., 2021). Upon detection of BAC-UPP complexes, SapAB switches its role from repressor to activator of SapS, resulting in kinase autophosphorylation (Koh *et al*., 2021). SapS can then phosphorylate SapR, which in turn induces the production of the resistance transporter RapAB. RapAB, again based on evidence from its *B. subtilis* homologue, frees UPP from the inhibitory grip of BAC using a target protection mechanism (Kobras *et al*., 2020), allowing dephosphorylation of UPP and continuation of cell wall synthesis, rendering the cell resistant to BAC. The equivalent system in *B. subtilis* is a self-contained module, involving only a single transporter (BceAB) and TCS (BceRS) (Ohki *et al*., 2003) with no known further genes involved in its signalling or resistance mechanism. However, in *E. faecalis*, we observed that the TCS SapRS is itself transcriptionally induced by BAC, however we had not identified the regulatory system that controls this expression (Gebhard *et al*., 2014).

Interestingly, recent work by the Arias lab demonstrated that when experimentally evolving *E. faecalis* for DAP resistance in a Δ*liaFSR* background, mutations were observed in the sensory transporter SapAB (Prater *et al*., 2021, 2019), suggesting a functional link between the Sap and Lia systems. In *B. subtilis*, the Lia system is known to be one of the main components of BAC resistance (Mascher *et al*., 2003; Rietkötter *et al*., 2008), but there is currently no evidence for a role of LiaFSR in response to BAC in *E. faecalis*. Neither is there any indication of a role for the Sap system in responding to DAP exposure. However, this recent evidence suggested an interplay between both the Sap and Lia systems and that, potentially, both systems may be contributing to DAP resistance.

In accordance with the need to deepen our knowledge of the CESR in *E. faecalis,* in this study we sought to examine the potential interplay between the Sap and Lia systems and investigate the involvement of further genes associated with this regulatory network. Using gene deletions, analysis of promoter activity and mutation of putative regulator binding sites, we provide evidence of the activation of LiaFSR signalling in response to BAC and describe a direct functional link between the Lia and Sap systems by demonstrating the regulation of *sapRS* by LiaR in response to antibiotic exposure. We further identify the *dltXABCD* operon, a known determinant of DAP resistance (Prater *et al*., 2019), as a target of SapRS regulation. Through a detailed analysis of promoter activation in response to DAP exposure, we show that the same regulatory network also controls the response of *E. faecalis* to this antibiotic. The complex regulatory architecture of LiaFSR controlling expression of *sapRS*, which in turn controls a set of BAC and DAP resistance genes appears unique to enterococci. It may allow sensitive and robust changes in gene expression through ‘priming’ of the SapRS response via LiaFSR-dependent upregulation of SapRS levels and offers an explanation why mutations in DAP-resistant clinical isolates and experimentally evolved strains so often are found in these regulatory genes.

## Results

### LiaFSR responds to bacitracin and controls expression of *sapRS*

As mentioned above, from previous work we had identified increased expression of *sapRS* under exposure to BAC or mersacidin (Gebhard *et al*., 2014). Interestingly, we also had shown that *sapRS* was not autoregulated, in contrast to other two component systems such as CroRS and LiaFSR (Comenge *et al*., 2003; Jordan *et al*., 2006), and regulation did not depend on either SapAB or RapAB (Gebhard *et al*., 2014). We therefore here first aimed to identify the regulator of *sapRS*. For possible candidates, we considered potential regulators that are common amongst the Firmicutes and are known to respond to BAC. The most probable candidate appeared to be LiaFSR, often referred to as the ‘master regulator’ of the CESR in *Bacillus subtilis* (Mascher *et al*., 2003) and recently shown to have a possible functional link to the Sap system (Prater *et al*., 2021, 2019).

To test if LiaFSR might regulate *sapRS* expression, we first had to establish if LiaFSR was able to respond to BAC treatment. It is well established that *liaX* is under the control of LiaFSR (Khan *et al*., 2019), and therefore we exposed *E. faecalis* harbouring a P*_liaX_*-*lacZ* transcriptional fusion to increasing levels of BAC, as a readout for LiaFSR activity. The results showed a ∼10-fold increase in P*_liaX_* activity at 32 μg ml^−1^ BAC compared with untreated cells (Fig. 1A, black line), indicating that LiaFSR can indeed respond to BAC exposure in *E. faecalis* albeit weakly. Addition of higher concentrations of BAC significantly impacted cellular growth and was therefore not used to test promoter induction.

**Figure 1.**
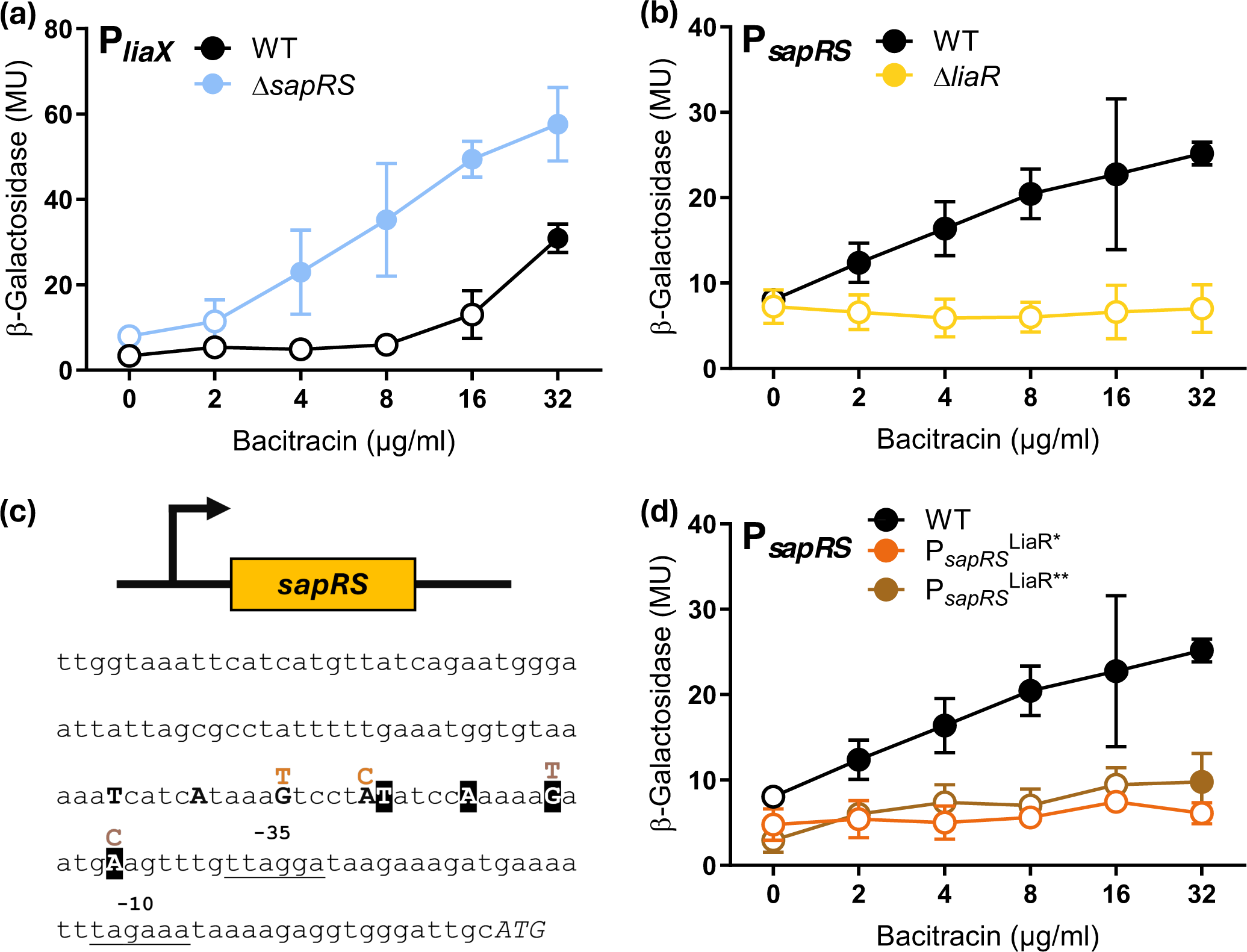
The LiaFSR system responds to bacitracin exposure and induces the expression of both *liaXYZ* and *sapRS*. Cells of *E. faecalis* JH2-2 wild type or isogenic deletion strains harbouring a P*_liaX_-lacZ* (a) or P*_sapRS_-lacZ* (b) transcriptional fusion were grown to exponential phase and challenged with increasing concentrations of bacitracin. b-galactosidase activity, expressed as Miller units (MU), was assayed following 1h incubation. (c) Schematic of the promoter region upstream of *sapR*. The start codon is shown as ATG and putative -10 and -35 promoter elements are underlined. The two proposed LiaR binding sites are indicated in bold capitals, black on white, and white on black, respectively. Mutations introduced in the first and second binding site are shown in orange and brown, respectively. (d) P*_sapRS_-lacZ* transcriptional fusions carrying the wild-type promoter sequence or the mutations in the first (LiaR*) or second (LiaR**) binding site of the promoter were introduced into wild-type *E. faecalis* and assayed as before. The colour code corresponds to that used in panel c. Please note the data for the native promoter is identical between panels b and d. Results are means and standard deviations for three biological repeats. The significance of induction relative to untreated cells was calculated for each strain by a two-way ANOVA test. Significance is indicated by a filled symbol (p < 0.05); unfilled symbols represent no significant increases compared to uninduced conditions. Numerical data for these graphs is provided in Table S4.

Following this, we tested if LiaR could regulate the expression of *sapRS*. To do this, we monitored the response of a transcriptional P*_sapRS_*-*lacZ* fusion to increasing BAC. The results showed that in wild-type *E. faecalis* carrying the transcriptional *sapRS-lacZ* fusion, BAC concentrations of 16 µg ml^-1^ or higher led to significant induction of the reporter, with a three-fold increase compared to uninduced cells at 32 µg ml^-1^ (Fig. 1B, black line). Deletion of *liaR* in the reporter strain resulted in a complete loss of *sapRS* induction, with expression remaining at basal levels (Fig. 1B, yellow line). This indicated that LiaR indeed regulates the expression of *sapRS* in response to BAC exposure, presenting first evidence of a direct functional link between the Lia and Sap regulatory systems.

### The bacitracin-response of LiaFSR is masked by the primary resistance determinant, RapAB

A surprising feature of the BAC-induced LiaR activity was the high concentration required to trigger a response, as *liaX* expression only significantly increased at 32 µg ml^-1^. One possible explanation for this might be that the response was masked by other components of the BAC stress response in *E. faecalis*, for example RapAB, which plays an active role in the removal of BAC from UPP (Gebhard *et al*., 2014), protecting the cell from damage. In *B. subtilis*, it was observed that the RapAB equivalent, BceAB, was the primary response to protect the cell from BAC exposure and masked the response of the Lia system (Radeck *et al*., 2016). To prevent the induction of the SapRS-dependent BAC resistance genes of *E. faecalis*, including *rapAB* (Gebhard *et al*., 2014), and thus remove potential interference with Lia signalling, we therefore introduced the P*_liaX_*-*lacZ* transcriptional fusion into the Δ*sapRS* background. Compared to the wild type, the expression of the *liaX* promoter in response to BAC challenge of the deletion strain showed increased sensitivity, significantly inducing expression from 4 μg ml^−1^ BAC and reaching overall higher activities (Fig. 1A, blue line). This showed that the weak response of Lia signalling to BAC in the wild type was indeed due to masking effects of the resistance genes controlled by SapRS, implying the presence of multiple layers of protection, similar to those observed in *B. subtilis* (Radeck *et al*., 2016). These data also provided further indication of physiological links between the SapRS- and LiaFSR-dependent components of the CESR in *E. faecalis*.

To test the physiological implications of LiaR controlling *sapRS* expression, the deletion strains were tested for their minimal inhibitory concentration (MIC) of BAC (Table 1). Consistent with our previous results (Gebhard *et al*., 2014), deletion of the known BAC resistance operon *rapAB* and the regulatory operons *sapRS* and *sapAB* led to a two- to four-fold decrease in MIC. Deletion of *liaR* only had a minor, if any, effect on MIC, with 8-16 µg ml^-1^ BAC required to prevent its growth, compared to 16 µg ml^-1^ for the wild type. This mild phenotype was consistent with LiaR acting as an inducer of the main BAC-responsive TCS, SapRS, rather than directly inducing the resistance genes. LiaR activity thus likely acts to amplify the response, but some resistance gene expression still occurs from basal levels of SapRS. This is addressed in more detail in the following section. Complementation of BAC sensitivity in the *liaR*-deleted strain with an ectopic copy of the gene driven by the native promoter of the *liaFSR* operon (i.e. a P*_liaF_-liaR* construct) unfortunately was unsuccessful. It had been shown previously for the LiaFSR system of *B. subtilis* that its function is exquisitely sensitive to the stoichiometry of the three proteins (Schrecke *et al*., 2013). It is therefore likely that a similar situation exists in *E. faecalis* and that we were not able to re-create the exact cellular level of LiaR needed for the regulatory system to function correctly. Complementation of reporter gene induction could not be tested, because all plasmids available for this study carry the same replication origin for *E. faecalis* and could therefore not be maintained in combination.

**Table 1.** Bacitracin and daptomycin sensitivity of *E. faecalis* strains.

| Strain or genotype | Minimal inhibitory concentration (MIC; $\mu\text{g ml}^{-1}$)<sup>a</sup> | |
| --- | --- | --- |
|  | Zn<sup>2+</sup>-Bacitracin | Ca<sup>2+</sup>-Daptomycin |
| JH2-2 | 16 | 16 |
| JH2-2 $\Delta\text{sapRS}$ | 4-8 | 4 |
| JH2-2 $\Delta\text{sapAB}$ | 4-8 | 4-8 |
| JH2-2 $\Delta\text{rapAB}$ | 4-8 | 8-16 |
| JH2-2 $\Delta liaR$ | 8-16 | 4-8 |
| JH2-2 $\Delta liaR/P_{liaF-liaR}$ | 8 | 4-8 |

<sup>a</sup> Results are from three independent repeat experiments. Where the MIC values differed between repeats, the range of obtained results is reported.

### A LiaR binding site is present in the *sapRS* promoter and required for induction

To confirm that the regulation of *sapRS* was directly due to LiaR activity, particularly in light of the lack of complementation of the resistance phenotype, we searched the promoter sequence upstream of the *sapR* start codon for a potential LiaR binding site. The promoter contained sequences with a good match to consensus -10 and -35 elements with conventional spacing (Fig. 1C, underlined). LiaR of *E. faecalis* has been shown to have a complex mechanism of promoter recognition, involving a primary consensus binding sequence and adjoining secondary recognition sequence, and activation involves a dimer to tetramer transition (Davlieva *et al*., 2015). This is further complicated by the consensus sequence having only four, evenly spaced, conserved residues: T-N_4_-C-N_4_-G-N_4_-A, with degeneracy observed between different LiaR targets (Davlieva *et al*., 2015). Searching the *sapR* promoter for a LiaR binding site revealed the existence of two such sequences, positioned directly next to each other and each with only a single mismatch to the consensus (Fig. 1C, bold and white-on-black letters). From this, it was not possible to predict which was the most likely candidate binding site, or if these two sequences might represent the previously described primary and secondary recognition sequences.

To test this experimentally, we generated two new P*_sapR_*-*lacZ* fusions by DNA synthesis, one in which two bases of the promoter-distal putative binding sites had been mutated, and one where two positions of the promoter-proximal binding site had been changed (Fig. 1C, coloured letters). When either of these constructs was introduced into wild-type *E. faecalis* and assayed for induction by BAC, no response could be detected at any tested antibiotic concentrations (Fig. 1D). These data strongly suggest that the identified sequences indeed constitute the LiaR binding site and are likely the pair of binding sites required for the primer-to-tetramer transition mechanism of LiaR-dependent promoter activation. This further confirmed direct regulation of *sapRS* expression by the LiaFSR system.

### SapRS controls the expression of the *dltABCD* operon

In *B. subtilis*, both the Lia and Bce systems act as self-contained modules, controlling the regulation of only a single resistance operon each, *liaIH* and *bceAB* respectively, each encoded adjacently to its regulatory operon on the chromosome (Ohki *et al*., 2003; Mascher *et al*., 2003; Wolf *et al*., 2010). However, the direct interaction between Lia and Sap signalling demonstrated above suggests that the regulatory setup in *E. faecalis* is much more complex. This is further supported by evidence from the literature, with the indication that the Lia system has a larger operon than just itself and *liaXYZ* (Khan *et al*., 2019). There is also a proposal that Lia contributes to DAP resistance through the regulation of *dltXABCD* (*dlt*), although this is likely an indirect effect also involving the *yvcRS* operon (Prater *et al*., 2021, 2019), which encodes the sensory transporter SapAB that works together with SapRS in signalling of BAC stress, as explained above. Evidence for the link between LiaFSR, SapAB-SapRS and the *dlt* operon comes from a study evolving *E. faecium* for DAP resistance in the absence of LiaFSR. Suppressor mutations occurred in the *sapAB* (*ycvRS*) operon and correlated with an increase in *dlt* transcription when measured by qPCR and an increase in cell surface charge (Prater *et al*., 2021, 2019). Dlt is responsible for the D-alanylation of teichoic acids on the bacterial cell surface, resulting in a decrease in the negative charge of the cell envelope (Peschel *et al*., 1999), a mechanism commonly involved in DAP resistance amongst the low-CG bacteria (Tran *et al*., 2015). Moreover, the *dltABCD* operon is located directly adjacent to *sapAB* on the chromosome, further supporting a functional link between the genes. To expand our understanding of the Lia/Sap regulatory network and how a BAC resistance system might be linked to DAP resistance, we therefore next sought to investigate the contribution of LiaFSR and SapRS to *dlt* regulation.

To this end, we first constructed a transcriptional P*_dltX_-lacZ* fusion to test *dlt* induction by BAC. The results showed that in wild-type JH2-2 carrying the fusion, increasing BAC concentrations led to a dose-dependent increase in *dlt* expression, resulting in ∼4-fold higher activity at 32 μg ml^−1^ compared to untreated conditions (Fig. 2A, black line), showing that *dlt* expression is induced in response to BAC in *E. faecalis*. When we tested the response of the reporter in strains carrying deletions of either *sapRS* or *sapAB,* the results showed a decrease in basal activity in both strains and a complete loss of the promoter’s BAC response in Δ*sapR* (Fig. 3A, blue and magenta lines). Loss of SapAB still resulted in induction compared with uninduced cells, but overall activities were considerably lower than in the wild-type strain. This indicated that SapRS was essential for *dlt* expression in response to BAC, but a residual amount of *dlt* induction remained in the *sapAB* deletion. This is consistent with the less direct regulatory role of SapAB, i.e. via controlling SapRS signalling activity and not the target genes directly. While SapAB is required to activate SapRS signalling, it does not influence the expression of *sapRS* (Gebhard *et al*., 2014), which is instead controlled by LiaFSR as we have shown above. Thus, in a *sapAB-*deleted strain, the cellular amount of SapRS will still increase in response to BAC due to LiaR activity, leading to higher basal activity of the TCS, which can explain the weak induction of P*_dltX_* -*lacZ* observed in this strain.

**Figure 2.**
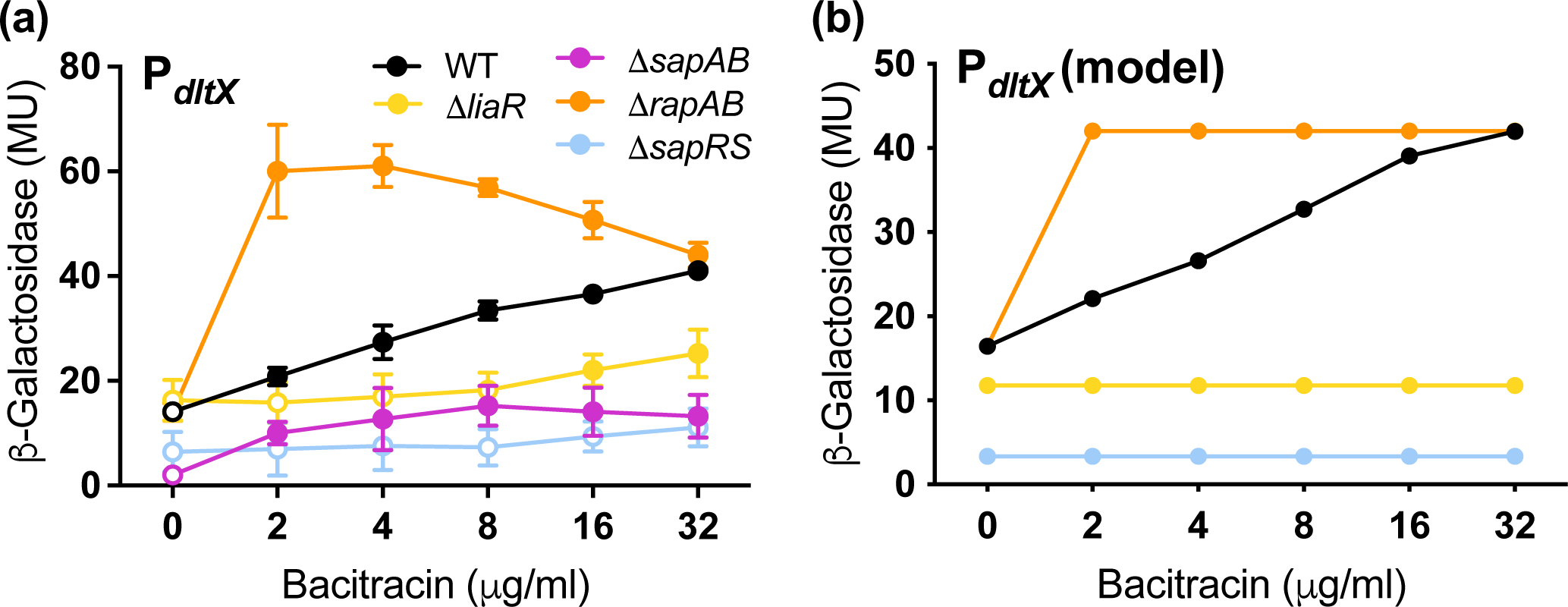
Induction of the resistance operon *dltXABCD* by bacitracin. Panel A indicates the experimental induction of *dltX.* Cells of *E. faecalis* harbouring a *PdltX-lacZ* transcriptional fusion were grown to exponential phase and challenged with increasing concentrations of bacitracin. Βeta-galactosidase activity, expressed as Miller units (MU), was assayed following 1h incubation in wild type (WT) and deletion strain backgrounds. Results are means and standard deviations for three biological repeats The significance of induction relative to untreated cells was calculated for each strain by a one-tailed, pairwise *t*-test analysis. Significance is indicated by a filled symbol (*p* < 0.05), unfilled symbols represent no significant increases compared to uninduced conditions. Panel B indicates the mathematical model of *dltX* induction in the *E. faecalis* strains indicated in panel A. Numerical data for these results is provided in Table S4.

**Figure 3.**
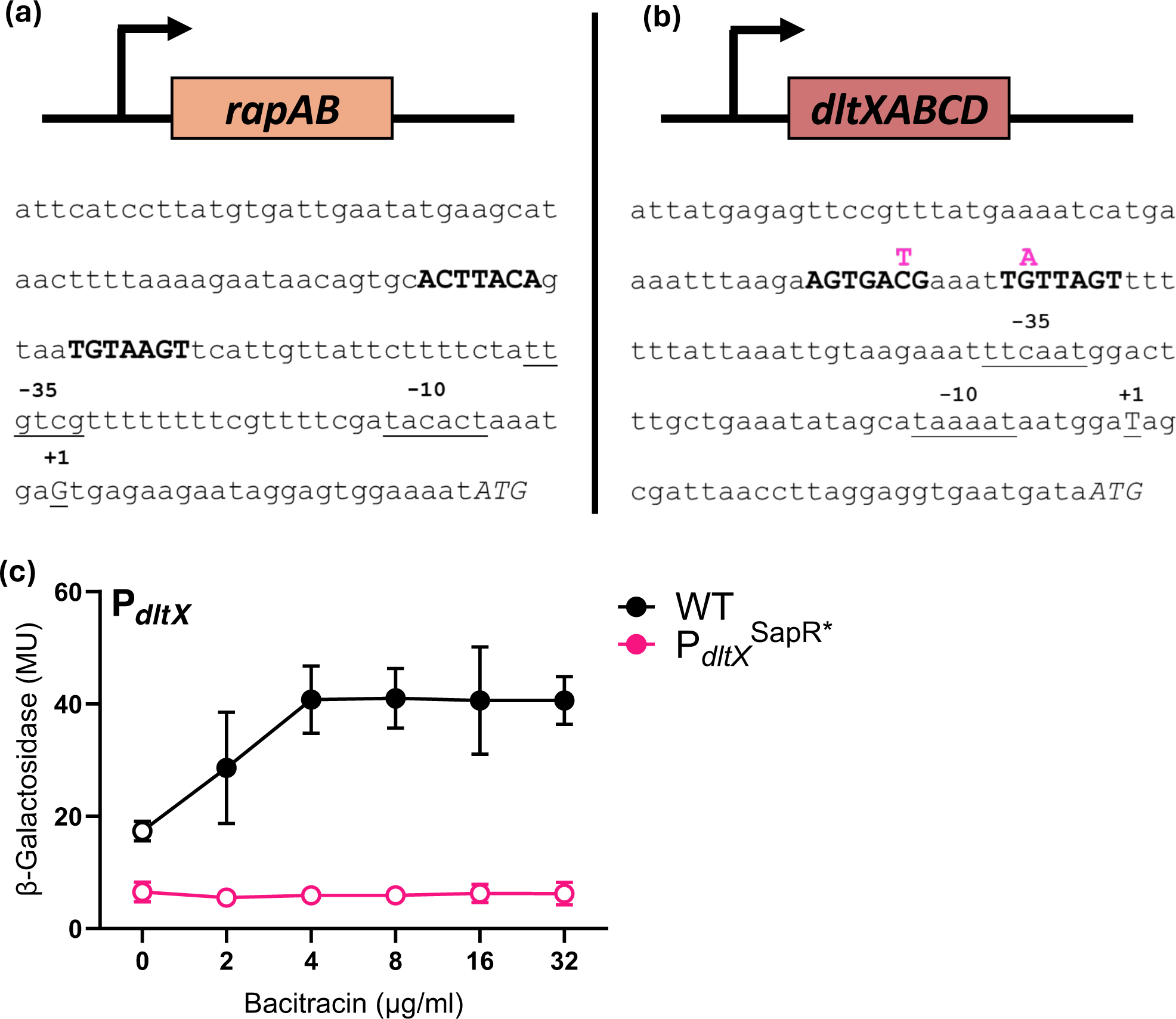
A putative SapR binding site in the *dltX* promoter is required for bacitracin-responsive promoter induction. (a, b) Schematics of the promoter regions of the known SapR target *rapAB* (a) and the newly identified target *dltXABCD* (b). The start codons are shown as ATG, and putative -10 and -35 promoter elements and likely +1 position are underlined. The known SapR binding site in P*_rapA_* and the proposed site in P*_dltX_* are shown in bold. Mutations introduced in the proposed binding site are shown in pink. (c) The P*dltX-lacZ* transcriptional reporter and a derivative carrying the two mutations in the putative SapR binding site from panel b were introduced into wild-type *E. faecalis.* Exponentially growing cells were challenged with increasing concentrations of bacitracin. β-galactosidase activity, expressed as Miller units (MU), was assayed following 1h incubation. Results are means and standard deviations for three biological repeats. The significance of induction relative to untreated cells was calculated for each strain by a two-way ANOVA test. Significance is indicated by a filled symbol (*p < 0.05*); unfilled symbols represent no significant increases compared to uninduced conditions. Numerical data for these graphs is provided in Table S4.

Overall, the behaviour of the *dltX* promoter was very similar to that of the previously identified SapR target, the *rapAB* operon (Gebhard *et al*., 2014). A P*_rapA_-lacZ* fusion responded to BAC challenge with a five-fold induction of activity (from 23 to 112 Miller Units), while this activity was reduced by 80% in a *sapS*-deleted strain, and low expression but observable BAC-induction was observed in the *sapAB* deletion background (Gebhard *et al*., 2014). The similar behaviour of the two promoters was consistent with both being controlled by the same regulator.

### A SapR binding site is present in the *dltXABCD* promoter and required for induction

To confirm that *dlt* operon control was due to direct activation by SapRS, we again inspected the sequence of the promoter region upstream of the first gene of the operon, *dltX*. This revealed a putative SapR binding site, similar to the one we had previously identified in the *rapAB* promoter (Gebhard *et al*., 2014) and in agreement with the consensus sequence for BceR-like regulators in Firmicutes bacteria, i.e. an inverted repeat around a central ACA-N_4_-TGT motif (Dintner *et al*., 2011) (Fig. 3). We next constructed a reporter gene fusion where the putative SapR binding site was mutated at one position in each half of the core motif (Fig. 3B, coloured letters). When wild-type *E. faecalis* carrying this reporter construct was challenged with BAC, induction was completely abolished (Fig. 3C), showing that the putative binding site was indeed essential for promoter activation and that control of *dlt* operon expression was likely a direct activity of SapR.

Because we had shown above that LiaR regulates *sapRS* expression, we next aimed to find out if deletion of *liaR* also influenced *dlt* expression. When we tested the response of the *dltX* promoter fusion to BAC in a strain carrying deletion of *liaR*, we observed that deletion of *liaR* resulted in a noticeably weaker amplitude of *dlt* expression in response to BAC, being ∼2-fold lower than the wild-type strain at 32 μg ml^−1^ (Fig. 2A, yellow line). In addition, the sensitivity of the *dlt* promoter response was much lower in the *liaR* deletion compared with the wild-type response, with significant activation over baseline occurring at 16 μg ml^−1^ rather than at 2 μg ml^−1^ in the wild type. However, there was no effect on basal activity of *dlt*, in contrast to deletion of *sapR* and *sapAB*. These data are consistent with LiaR regulating *sapRS* expression, and SapRS being the actual regulator of *dlt* expression in response to BAC.

### The primary bacitracin resistance determinant, RapAB, provides a negative feedback on the wider regulatory network

As we had observed before with the *liaX* and *sapRS* promoters, we also wanted to investigate if the protection provided by the SapRS-target RapAB was dampening the expression of *dlt* in response to BAC. To examine this, we tested the response of *dlt* expression in the absence of *rapAB.* In a *ΔrapAB* background, the *dlt* promoter demonstrated markedly increased sensitivity, and a much stronger response to BAC, resulting in a ∼3-fold increase in expression at 2 μg ml^−1^ when compared to the wild-type (Fig. 2A, orange line). This response demonstrated the presence of a layered protection with RapAB activity moderating *dlt* expression.

Overall, these findings were rather surprising, as the signalling pathway appeared remarkably complex to result in a relatively simple outcome, i.e. inducing the expression of two resistance genes in response to an antibiotic. To test if our understanding of the regulatory pathway was plausible, we therefore developed a representative mathematical model to see if this would reproduce the behaviour we had observed in the experiments. At the core, this model was based on a simplified form of the flux-sensing mechanism described previously for the *B. subtilis* BceRS-BceAB system (Fritz *et al*., 2015), which was then expanded upon to reproduce the hypothesised network structure investigated here. In brief, the model considered BAC binding to UPP to form BAC-UPP complexes with the on-rate dependent on the BAC concentration. These complexes then drive transport activity of, in this case, SapAB, according to a soft-switch type functional response (see Methods), which the model translates into activation of SapRS and thus *dlt* expression. Importantly, SapRS activity also drives expression of the resistance transporter operon, in this case *rapAB*, and production of RapAB leads to a reduction in formation of UPP-BAC complexes due to the target protection activity of the transporter (Fritz *et al*., 2015; Kobras *et al*., 2020). This creates the negative feedback loop that is characteristic of the flux-sensing mechanism and leads to the gradual response behaviour of the output promoters (Fig. 2B, black symbols). To adapt this model to the Lia-Sap regulatory pathway of *E. faecalis*, we additionally considered the activity of the Lia system. This was modelled to also respond to BAC-UPP complexes in a switch like manner representing the generation of cellular damage caused by these complexes. The model then linked the Lia and Sap components by modulating the SapRS signalling output (i.e. *dlt* and *rapAB* expression) according to Lia activity, driven by BAC. The mathematical details of the model are explained in the Methods section. Fitting to experimental data was performed by matching differential activity of SapRS against both *rapAB* and *dlt* recorded at different BAC concentrations.

This model accurately reflected the behaviour of our experimental strains, depicting the same gradual response to BAC of the *dltX* promoter in wild-type *E. faecalis* (Fig. 2B, black symbols), as well as the hypersensitive response in the *rapAB* deletion strain, where the negative feedback from RapAB-driven removal of BAC was missing (orange symbols). Importantly, the model gave the same complete loss of *dltX* activity when *sapRS* was deleted (blue symbols) as observed experimentally, as well as the normal basal level activity but loss of BAC-dependent induction when *liaR* was deleted (yellow symbols). To simplify the previous comprehensive model of signalling by Bce-like systems, we here considered the sensory complex of SapAB and SapRS as a single agent in the network. While this allowed us to test the new connection to Lia-dependent signalling and the additional target operon *dlt* without overly complicating the model, the model now does not differentiate between SapAB and SapRS activities separately. Therefore, the *sapAB* deletion strain from the experimental data was not specifically considered in the theoretical analyses.

Overall, the close agreement between theoretical and experimental data strongly suggested that our reconstruction of the regulatory pathway and connection between the Lia and Sap systems was theoretically plausible and that no further major players were needed to explain the behaviour of the *dltX* target promoter in the various mutant backgrounds analysed here.

### The LiaFSR-SapAB-SapRS network responds to daptomycin challenge

We had now established a sequential order of *dlt* regulatory control: in response to BAC, LiaFSR induces the expression of *sapRS*, and in turn SapRS, activated by its sensory transporter SapAB (Gebhard *et al*., 2014), induces the expression of *dlt*. As described in the introduction, the Lia system of *E. faecalis* is primarily known for its contribution to DAP resistance in clinical strains (Arias *et al*., 2011), and as we showed above, only makes a minor contribution to BAC resistance. Moreover, increased expression of *dlt* is a known cause of DAP resistance (Prater *et al*., 2019). It was therefore surprising to find that *dlt* regulation was separated from LiaFSR activity by the BAC-responsive SapAB-SapRS regulatory system. We were particularly intrigued by this network structure, because the mode of action of DAP makes it very unlikely to be recognised by SapAB, and thus DAP exposure should not trigger SapRS activation.

To dissect the expected differences in sensory perception of the two antibiotics, their mode of action needs to be considered. BAC forms a complex with UPP, which acts as the input for SapRS signalling via SapAB (Kobras *et al*., 2020); at the same time, BAC induces cell envelope damage, triggering LiaFSR activation (Wolf *et al*., 2012). DAP, on the other hand, forms a tripartite complex with Lipid II and phosphatidyl-glycerol in the membrane (Grein *et al*., 2020), which leads to dissociation of components of the cell wall biosynthetic machinery from the membrane and thus inhibits cell wall synthesis (Müller *et al*., 2016). Eventually, DAP also causes membrane depolarisation (Müller *et al*., 2016), although it has not yet been shown if this is also a direct consequence of the tripartite complex formation. The DAP-induced cell envelope damage activates the LiaFSR response (Hachmann *et al*., 2009; Wecke *et al*., 2009). However, because DAP does not bind to UPP or the sugar-pyrophosphate moiety of Lipid II, which is the common feature of substrates for BceAB-type transporters (George *et al*., 2022; Kobras *et al*., 2020), it is not expected to be recognised by SapAB to trigger SapRS signalling. To understand the rationale of directly linking Lia and Sap signalling in *E. faecalis* to control a known DAP-resistance operon (*dlt*), we therefore set out to investigate how each of the network nodes responds to this second antibiotic.

The first step in the signalling cascade is the activation of P*_sapR_* by LiaFSR signalling. When we exposed wild-type *E. faecalis* carrying the *P_sapRS_-lacZ* transcriptional fusion to increasing concentrations of DAP, a gradual activation was observed from 1 μg ml^-1^ to an approximately four-fold maximal activation at concentrations above 8 μg ml^-1^ of the antibiotic (Fig. 4A, black line). Consistent with LiaR being the sole regulator of the *sapR* promoter, deletion of *liaR* completely abolished promoter induction, while deletions of *sapRS* or *sapAB* had no effect (Fig. 4A, coloured lines).

**Figure 4.**
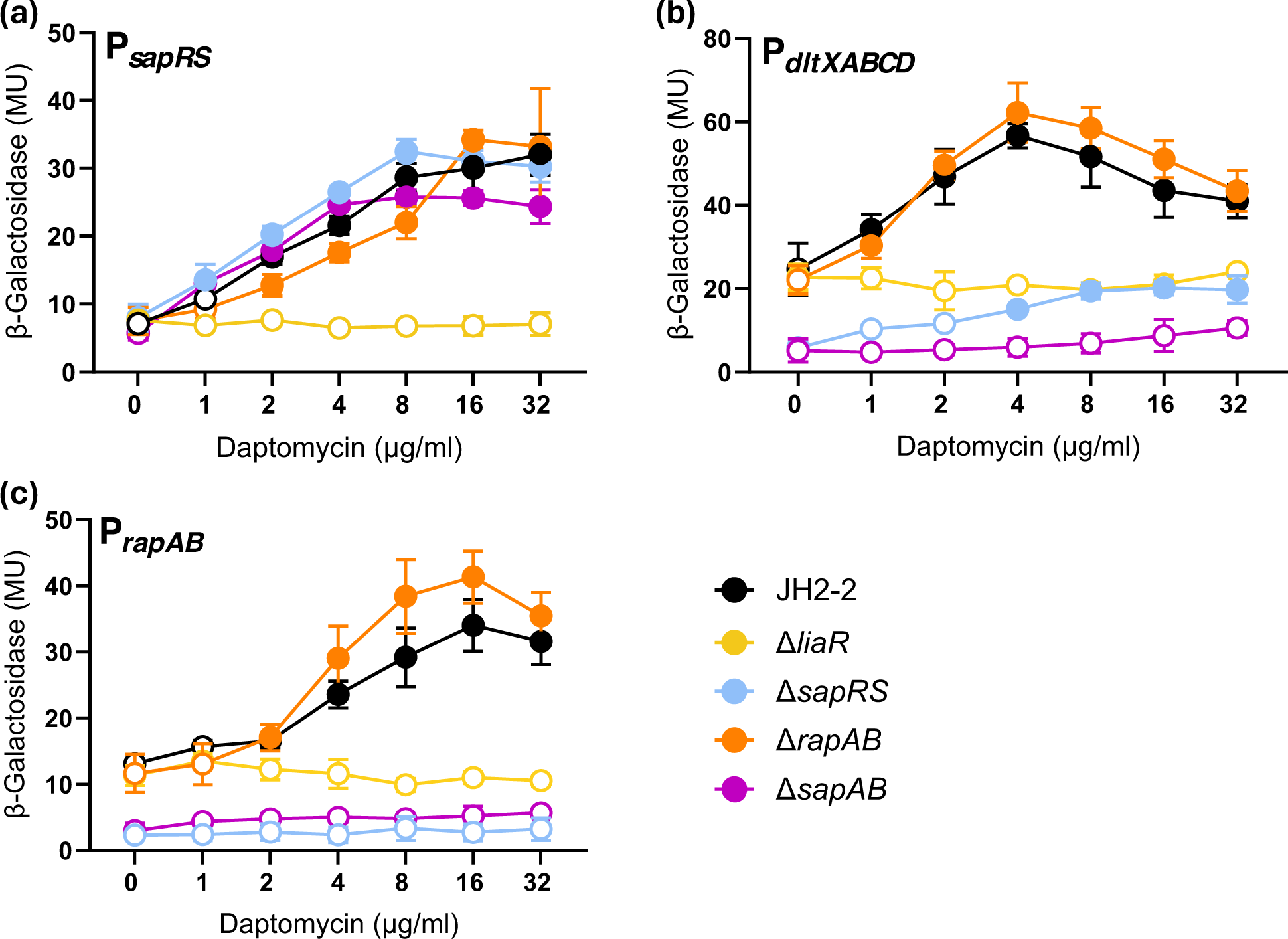
The Daptomycin-response of the regulatory network. *E. faecalis* cells harbouring *P_sapRS_-lacZ* (a), *P_dltX_-lacZ* (b) or *P_rapAB_-lacZ* (c) transcriptional fusions were grown to exponential phase and challenged with increasing concentrations of daptomycin. b-galactosidase activity, expressed as Miller units (MU), was assayed following 30 minutes incubation in wild type (WT) or deletion strain backgrounds, as indicated. Results are means and standard deviations for three biological repeats. The significance of induction relative to untreated cells was calculated for each strain by a two-way ANOVA test. Significance is indicated by a filled symbol (P < 0.05), unfilled symbols represent no significant increases compared to uninduced conditions. Numerical data for these graphs is provided in Table S4.

Next, we tested the response of the P*_dltX_-lacZ* transcriptional fusion, identified as a SapRS target above, to increasing DAP concentrations. This reporter also showed a dose-dependent response to the antibiotic from 1 μg ml^-1^, with an approximately three-fold maximal activation observed at 4 μg ml^-1^ DAP (Fig. 4B, black line). Similarly to the BAC response, and consistent with the direct regulatory control of the promoter by the SapAB-SapRS system, loss of either of their encoding operons reduced the basal activity of the *dlt* promoter and abolished or greatly reduced the DAP response (Fig. 4B, light blue and purple lines). The slight remaining induction in the *sapRS*-deleted strain is discussed further below. Deletion of *liaR* only abolished the DAP response but did not affect the basal activity (Fig. 4B, yellow line), consistent with the corresponding BAC-data and the indirect effect of Lia signalling on *dlt* operon expression. Overall, these data closely reflected the behaviour of the regulatory network in response to BAC, with the exception of the lack of negative feedback provided by RapAB, which is addressed below. These results also provided experimental proof that the Sap signalling system was indeed a component of the Lia-dependent control of DAP resistance gene expression in *E. faecalis*, even though DAP did not belong to its expected substrate spectrum.

In this light, it was intriguing that deletion of *sapAB* abolished induction of the *dlt* promoter. Bce-type transporters, such as SapAB, generally activate signalling of their cognate two-component system in the presence of a suitable substrate (Staroń *et al*., 2011; Gebhard *et al*., 2014; Gebhard and Mascher, 2011). We had therefore considered that SapAB might be dispensable for DAP signalling. Recent data from cryo-EM structures of the BceAB-BceS complex from *B. subtilis*, however, had indicated that transporter and histidine kinase can mutually affect each other’s activity (George and Orlando, 2023), which may suggest that in *E. faecalis* the physical presence of SapAB is required for optimal SapS activity, even in the absence of a substrate. Another curious observation in the data was the slight DAP response, albeit at a much-reduced level, which was still apparent in the *sapRS* deletion background (Fig. 4B, light blue line). This was entirely unexpected, if SapR was indeed the sole regulator of *dltX*, and is also in contrast to the BAC response of the promoter, where no induction remained after deletion of *sapRS*. These data suggest that yet another regulatory pathway in the cell, one that responds to DAP but not BAC, contributes to the control of *dlt* operon expression.

Earlier we had shown that *rapAB* deletion had a negative effect on the BAC response, resulting from the removal of the antibiotic and therefore the stimulus from the cell. As DAP is not considered a suitable substrate for BceAB-family transporters, it was not expected that RapAB could remove DAP. Consistent with this, no effect on regulation of either the LiaR-target P*_sapR_*, or the SapR target P*_dltX_* was observed in the *rapAB*-deleted strain (Fig. 4A&B, orange lines). Nevertheless, the regulatory network structure described here places the *rapAB* operon, which is a known direct target of SapR (Gebhard *et al*., 2014), also under control of a DAP-responsive signalling pathway. We therefore tested a previously constructed P*_rapA_-lacZ* reporter construct (Gebhard *et al*., 2014) for its response to DAP challenge. Indeed, the results showed that the *rapA* promoter responded with an approximately three-fold induction to DAP, and the effects of the different gene deletions were very similar to what we had observed for the *dlt* promoter (Fig. 4C). The response amplitude was lower than the five-fold induction reported previously for BAC challenge of the same reporter (Gebhard et al, 2014), which can likely be explained by the amplifying effect of SapAB-SapRS signalling with BAC, but not DAP. Importantly, no partial induction of P*_rapA_* was observed in the *sapRS* deleted strain, suggesting that the additional, as yet unidentified, regulatory control of the *dlt* operon implied by the data shown in Fig. 4B must be specific to that target promoter.

### The regulatory network contributes to DAP resistance of *E. faecalis*

Finally, to test if all regulatory components are required to protect *E. faecalis* against DAP, the deletion mutants were tested for their MIC of the antibiotic. For all deletions of regulatory components, a two- to four-fold reduction in MIC from 16 mg ml^-1^ in the wild type to 4-8 mg ml^-1^ in the deletion strains was observed (Table 1). This was consistent with their contribution to *dlt* operon expression. While for BAC resistance, LiaR played a lesser role, for DAP resistance no such difference could be detected between the Lia and Sap components of the network. This is consistent with the BAC response involving an active amplification of signalling by the SapAB-SapRS component, whereas the DAP response cannot be amplified due to lack of substrate recognition by the transporter. Also consistent with the transcriptional reporter data and inability of BceAB-type transporters to recognise DAP, loss of RapAB had no significant effect on DAP resistance (Table 1). Unfortunately, despite numerous attempts, we could not construct a *dlt* operon deletion without causing significant polar effects. Therefore, the precise contribution of the Dlt system to DAP resistance, e.g. relative to other LiaFSR targets, could not be determined here.

## Discussion

In this study, we aimed to expand our understanding of the response of *E. faecalis* to cell envelope targeting antibiotics and investigate a potential functional link between the damage-sensing LiaFSR signalling pathway and the BAC-responsive SapAB-SapRS system. Our findings revealed the transcriptional regulation of the BceRS-type TCS SapRS by LiaFSR. In addition, we also demonstrated the regulation of the DAP resistance operon *dltXABCD* by SapRS. Even though the Sap system had been shown previously to control BAC resistance and possess a very narrow inducer range (Gebhard *et al*., 2014), our results showed that the regulatory interplay between Lia and Sap signalling allows the entire network to respond to both BAC and DAP challenge. Susceptibility testing showed that within this network, protection against BAC was primarily provided by the SapRS-controlled transporter RapAB, while DAP resistance required the full complement of regulatory components. This work therefore offers an explanation for previous observations of suppressor mutations in *sapAB* when a LiaR-deficient strain of *E. faecalis* was evolved for DAP resistance (Prater *et al*., 2021, 2019). A working model for the proposed regulatory network structure is presented in Figure 5.

**Figure 5.**
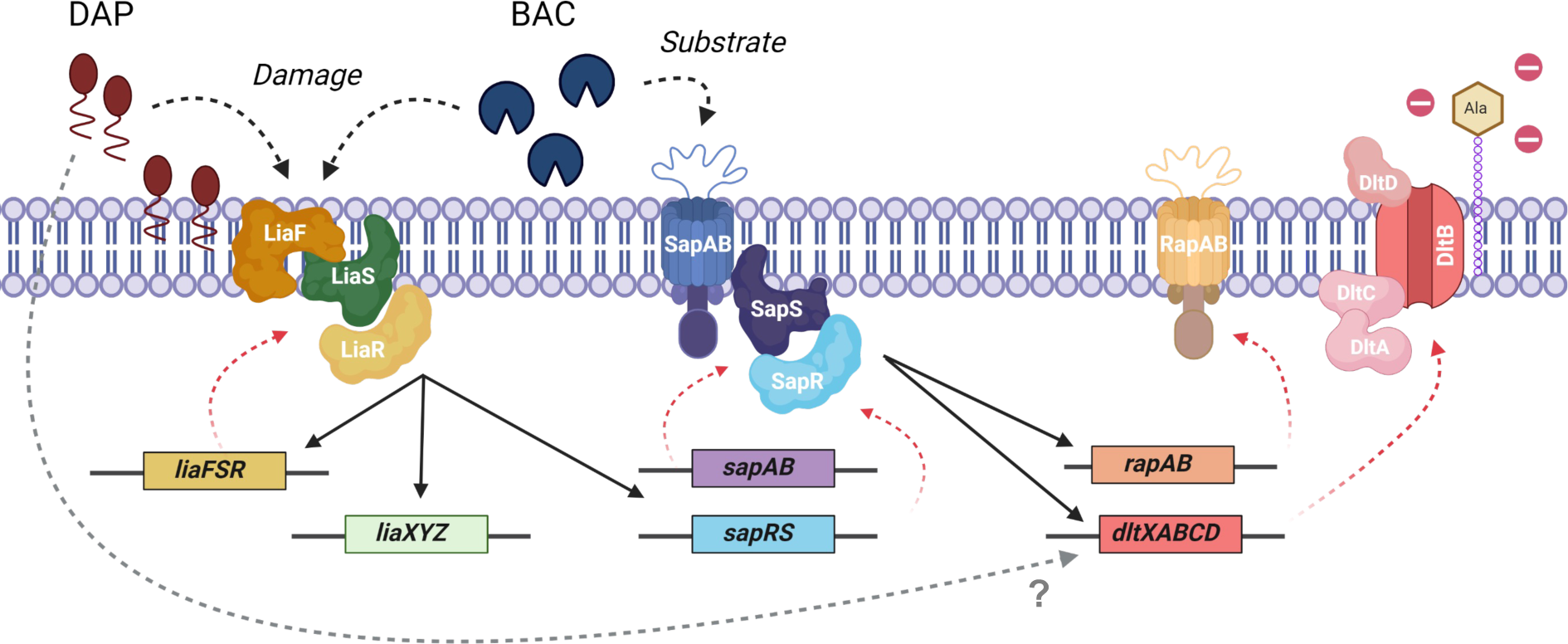
Proposed model of the antibiotic resistance network. Schematic illustrating the genes and proteins involved in antibiotic detection and resistance. Transcriptional regulation of genes is shown by black straight arrows, translation of genes to proteins is shown by dashed red arrows. Exposure to bacitracin (BAC) causes damage to the cell envelope, which acts as the stimulus for the removal of repression by LiaF, triggering the activation of LiaS. LiaS activity leads to phosphorylation of LiaR, which in turn induces the expression of its target promoters: *liaFSR*, *liaXYZ* and *sapRS*. Simultaneously, BAC is also detected by the SapAB/SapS sensory complex, leading to an amplification of the signalling response and via SapR. SapR then induces the expression of the resistance genes, *rapAB* and *dltXABCD,* which remove BAC from UPP and reduce the positive charge of the cell envelope, respectively. Conversely, exposure to daptomycin (DAP) again results in cell envelope damage, which triggers the phosphorylation of LiaR and induces the expression of *sapRS*. DAP is not recognised by SapAB and no amplification of the signal occurs. The LiaR-induced increase in SapRS production is nevertheless sufficient to increase basal SapRS activity, again leading to the induction of *rapAB* and *dltXABCD*. Of these, only the Dlt system is thought to counteract DAP action. The observed DAP-responsive increase of P*_dltX_* activity in the absence of SapRS by an as-yet-unidentified additional regulatory system is indicated by a grey, dashed arrow and a question mark.

LiaFSR is well-known to control DAP resistance in *E. faecalis*, and mutations that lead to constitutive activation of Lia signalling are commonly observed in resistant clinical isolates (Arias *et al*., 2011; Miller *et al*., 2013). While evidence is available that *dlt* operon over-expression leads to DAP resistance (Prater *et al*., 2019), work to date has mostly focussed on the primary LiaFSR target, the *liaXYZ* operon (Davlieva *et al*., 2015). The mechanism of resistance provided by this operon is not yet fully understood, but LiaX is thought to directly counteract the antibiotic and somehow mitigate its effects on the cytoplasmic membrane (Khan *et al*., 2019). Of the two known resistance determinants, one therefore is a direct target of LiaFSR signalling (*liaXYZ*), while the other (*dltXABCD*) is regulated via an expanded network involving the SapAB-SapRS system, as shown here. The reasons for this are not yet clear.

The situation is further complicated by our observation that some induction of *dlt* expression in response to DAP exposure still occurred in the *sapRS*-deleted strain (Fig. 4B), suggesting that yet another, so far unidentified regulator contributes to *dlt* regulation. Importantly, this response was specific to DAP, because no residual activation after BAC exposure was observed in the same genetic background (Fig. 2A). In *B. subtilis*, extracytoplasmic function (ECF) sigma factors play a crucial role in the cell envelope stress response, including control of the *dlt* operon (Helmann, 2016). In *E. faecalis,* to our knowledge, to date no ECF has been shown to be involved, although at least two putative ECF sigma factor/anti-sigma factor pairs are present in the genome, with homology to SigV and YlaC in *B. subtilis* (Casas-Pastor *et al*., 2021). SigV of *E. faecalis* has been shown to contribute to lysozyme resistance (Le Jeune *et al*., 2010) While SigV does not seem to control the *dlt* operon, at least in response to lysozyme (Le Jeune *et al*., 2010), further investigations are needed to explore the contribution of this sigma factor and YlaC to the extended cell envelope stress response network in this organism.

The second target of the Lia-Sap signalling pathway is the operon for the BAC resistance transporter RapAB. Consistent with our current understanding of transporters of the BceAB-type, RapAB could protect the cell from BAC, but not from DAP, as seen in the MIC values (Table 1). BceAB-type transporters recognise antimicrobial peptides that bind to either UPP or the UPP-sugar moiety of lipid II (George and Orlando, 2023; Kobras *et al*., 2020). DAP binds in a fundamentally different manner, forming a tripartite complex with lipid II and phosphatidyl-glycerol (Grein *et al*., 2020), and thus does not match the substrate spectrum of a transporter like RapAB. In close agreement with the susceptibility data, we could also show that the target protection mechanism, by which RapAB removes BAC from the cell, results in a negative feedback effect on the network’s response to BAC, whereas no such effect was observed during DAP exposure (Figs. 2 and 4). This raises the question why the *rapAB* operon was placed under control of a regulatory network that responds to DAP challenge, and indeed we could show that the *rapA* promoter itself is also significantly induced by this non-substrate antibiotic (Fig. 4). One explanation could be that this is simply an unintended consequence of the network structure, which may have primarily evolved to respond to antibiotics from which RapAB can protect. Alternatively, the answer may be found in the natural habitats of *E. faecalis* with high densities of competing micro-organisms, e.g. the soil or animal gut. Here, it may be an effective protection strategy to use a regulatory network that can respond to multiple different input antibiotics and respond by increased expression of resistance genes that can protect from a range of antibiotic mode-of-actions.

The regulatory network we describe here provides such a structure: it responds to two inputs, BAC and DAP, and it controls two outputs, RapAB and DltXABCD production. As stated above, of these inputs, BAC is most effectively counteracted by RapAB, while DAP is counteracted by changes in surface charge resulting from Dlt activity. Particularly given that LiaFSR signalling can directly lead to the activation of another DAP resistance operon, *liaXYZ*, it is surprising that the *dlt* operon is not simply also a direct LiaR-target. And moreover, why is the Lia system brought into the network to provide DAP-responsiveness to SapAB-SapRS signalling when one of the main SapR targets, *rapAB*, cannot protect the cell from this antibiotic? While we cannot yet answer this question, a speculative explanation for the convergence of Lia and Sap signalling within the same network might be found if we consider *E. faecalis* in its natural habitat.

As mentioned above, in an environment with high population density and diversity combined with scarcity of nutrients, it is plausible to expect that multiple species capable of antimicrobial production would secrete their inhibitory substances to gain a competitive advantage. The Lia-Sap regulatory network might equip *E. faecalis* with the means of reacting to one antimicrobial in a way that primes it to respond to further drugs. In this scenario, LiaFSR is the first level of the response. This system responds to the broadest substrate range, because its stimulus is cell envelope damage rather than a specific compound. Its activity then primes the more drug-specific Sap system to respond. If the encountered drug is a substrate for this system (here exemplified by BAC), SapAB will amplify the response through increasing SapS activity (Dintner *et al*., 2014; Gebhard *et al*., 2014; George and Orlando, 2023). The resulting high levels of RapAB will remove the antibiotic and thus dampen both the Lia and Sap responses through the negative feedback, preventing excess activation (Fig. 2) (Fritz *et al*., 2015). If the encountered antibiotic is not a RapAB substrate (here exemplified by DAP), the network responds sensitively on all levels (Fig. 4). Up-regulation of *rapAB* in this case may be an accident, or it could prepare the cell for the arrival of a suitable substrate, in which case its resistance would be instantly available. The additional input from an as-yet-unidentified DAP-responsive system on *dlt* expression might provide a further means of tuning the network response to the balance of antibiotics encountered.

To test this proposal, data are required on how the network responds to mixtures of various inducers but also in mixed microbial communities *E. faecalis* might naturally encounter. While such a comprehensive investigation is beyond the scope of the present study, we have carried out initial tests studying the inhibition of *E. faecalis* and derived mutants by a set of known antibiotic producers. The details of these experiments are presented in Supplemental File S2. In brief, we grew the different *E. faecalis* strains in agar overlays on plates that had been pre-grown with spots of four antimicrobial producing strains: the subtilin producer *Bacillus subtilis* ATCC6633 (Banerjee and Hansen, 1988), BAC producer *Bacillus lichenformis* ATCC10716 (Johnson *et al*., 1945), nisin A producer *Lactococcus lactis* NZ9000 (Kuipers *et al*., 1998) and nisin P producer *Streptococcus gallolyticus* AB39 (Aldarhami *et al*., 2020). While we cannot confirm that each of these strains produced their expected antimicrobial under the chosen growth conditions, each producer caused the appearance of an inhibition zone in the *E. faecalis* overlay, showing that some antimicrobial compounds were indeed secreted. In these antagonism assays, the broad importance of the role played by LiaFSR in response to a diversity of antimicrobials was well demonstrated, because deletion of *liaR* resulted in increased sensitivity against all tested antimicrobial producers, except for *L. lactis* NZ9000. In contrast, components of the Sap system only contributed to protection from the BAC producer *B. lichenformis* ATCC10716 and the subtilin producer *B. subtilis* ATCC6633. The only antimicrobial producer against which neither of the genes appeared to give a protective effect was the nisin-A producer *L. lactis* NZ9000. However, as stated we cannot currently ascertain whether the antimicrobial produced under these conditions was indeed nisin A, or whether *E. faecalis* uses alternative resistance mechanisms against nisin A not controlled by either the Lia or Sap systems. Overall, it appears that the regulatory network studied here plays a key role in protecting *E. faecalis* from antimicrobial peptide producing bacteria that it would realistically encounter in its natural habitats, offering an explanation for the existence of such a complex regulatory strategy.

The regulatory wiring we have uncovered here between the LiaFSR and the Bce-type SapAB-SapRS systems, resulting in the control of the *dltXABCD* and *rapAB* resistance operons, is remarkably complex compared with other Firmicutes bacteria. For example, in *B. subtilis* LiaFSR controls its own regulon, comprised solely of *liaIH* and its own encoding genes (Wolf *et al*., 2010). In *S. aureus*, the LiaFSR homologue VraRS has been shown to regulate the expression of multiple genes involved in cell wall biosynthesis (Wu *et al*., 2020), but has not been observed to regulate a BceRS-type TCS. Bce-type systems in most Firmicutes bacteria either solely control their own transporter, as found with BceAB-RS in *B. subtilis* (Ohki *et al*., 2003), or can be involved with other Bce-type systems, such as the BraRS-VraDE (Hiron *et al*., 2011) and GraRS-VraFG (Meehl *et al*., 2007) systems in *S. aureus*. In some cases, Bce-type systems can also control additional genes, for example the GraRS TCS has also been shown to regulate the *dlt* operon (Kraus *et al*., 2008; Cheung *et al*., 2014; Ledger *et al*., 2022). However, to our knowledge there have been no prior reports of direct regulatory relationships that hardwire the Lia and Bce-type pathways together.

Overall, our findings contribute to an increasing understanding of the regulatory network protecting *E. faecalis* from cell envelope attack and provide insights into the regulatory cascade that is involved in controlling resistance gene expression in response to cell envelope targeting antibiotics. Our findings also provide valuable context in which to better understand the emergence of DAP resistance in clinical environments. Ultimately, a system-wide understanding of antimicrobial resistance regulation may lead to identification of an ‘Achilles’ heel’ within the network and identify new therapeutic targets.

## Experimental Procedures

### Bacterial strains and growth conditions

All bacterial strains and plasmids used in this study are listed in Table S1 in the supplementary material. *E. coli* MC1061 was used for cloning with pTCVlac, and strain DH5α was used for all other cloning. *E. coli*, *Bacillus licheniformis* and *Bacillus subtilis* were routinely grown in lysogeny broth (LB) at 37°C with agitation (200 rpm). *Lactococcus lactis* was grown routinely in M17 supplemented with 0.5% lactose at 30°C without agitation. *Enterococcus faecalis and Streptococcus gallolyticus were* grown routinely in brain heart infusion (BHI) at 37°C without agitation, with media for the latter being supplemented with 5% (v/v) foetal bovine calf serum. Solid media contained 15 g l^-1^ agar. Selective media contained chloramphenicol (10 μg ml^−1^ for *E. coli*; 15 μg ml^−1^ for *E. faecalis*), kanamycin (50 μg ml^−1^ for *E. coli*; 1000 μg ml^−1^ for *E. faecalis*), spectinomycin (100 μg ml^−1^ for *E. coli*; 500 μg ml^−1^ for *E. faecalis*). For blue-white screening, 5-bromo-4-chloro-3-indolyl-β-d-galactopyranoside (X-Gal) was used at 120 μg/ml. Bacitracin was supplied as the Zn^2+^ salt. All media for experiments with daptomycin were supplemented with 500 µM CaCl_2_.

*E. faecalis* was transformed by electroporation as previously described(Cruz-Rodz and Gilmore, 1990). *E. coli* was transformed by heat-shock of CaCl_2_ competent cells, followed by 1 hour recovery time (Hanahan and Glover, 1985). Growth was measured as optical density at 600 nm (OD_600_) on a the Biochrom™ Novaspec Pro Spectrophotometer using cuvettes with 1 cm light path length or in 96-well plates with 100 μL culture volumes on a Spark^®^Microplate reader (Tecan).

### Construction of plasmids and genetic techniques

All primer sequences used for cloning are listed in Table S2 in the supplementary material. Transcriptional promoter fusions to *lacZ* in *E. faecalis* were constructed in the vector pTCVlac (Poyart and Trieu-Cuot, 1997). All fragments were cloned via the EcoRI and BamHI sites of the vector. Additionally, vectors with the mutated binding sites for LiaR and SapR were synthesized and sub-cloned into pTCVLac by GeneArt (ThermoFisher Scientific Inc., Darmstadt). Constructs for unmarked deletions in *E. faecalis* were cloned into pLT06 (Thurlow *et al*., 2009). For each gene or operon to be deleted, 700- to 1000-bp located immediately before the start codon of the first gene (“up” fragment) and after the stop codon of the last gene (“down” fragment) were amplified. The primers were designed to create a 17- to 20-bp overlap between the PCR products (Table S2), facilitating the fusion of the fragments by PCR overlap extension (Ho *et al*., 1989) and were subsequently cloned into the NcoI and BamHI site of the vector pLT06.

Gene deletions were performed as previously described (Thurlow *et al*., 2009). Briefly, following transformation of the parent strain with the temperature-sensitive vector pLT06, overnight cultures were grown at 30° containing chloramphenicol and reinoculated 1:1000 into 10 mL BHI the next morning. Cells were then grown at 30° for 2.5 hours, followed by increasing to 42° for a further 2.5 hours to force single-site integration. Cells were then serially diluted onto BHI agar containing chloramphenicol and X-Gal and incubated at 42°. Blue colonies growing at 42°C were screened for the targeted integration using PCR with primers flanking the site of integration. Positive clones were then serially passaged for two days from overnight culture in BHI medium with no selection at 30° to allow a second site recombination event. Cultures were then serially diluted on to MM9-YEG agar (Kristich *et al*., 2007b) containing 10 mM *p*-chloro-phenylalanine for counter-selection and X-Gal at 37°C. The resulting white colonies were screened for the deletion of the target genes by PCR. All cloned constructs were checked for PCR fidelity by Sanger sequencing, and all created strains were verified by PCR using the primers given in Table S2.

### Antimicrobial susceptibility assays

For antibiotic susceptibility assays, minimum inhibitory concentrations (MICs) were determined by broth dilution assays. Colonies of *E. faecalis* were suspended in sterile Phosphate Buffered Saline (PBS) to 0.5 McFarland standard turbidity and diluted 1:1,000 in a total volume of 1.5 ml BHI medium. For assays with daptomycin, the medium was supplemented with 500 mM CaCl_2_. Of the inoculated medium, 100 ml aliquots were added to wells containing two-fold serial dilutions of the antibiotic. After 24 h incubation at 37°C, growth was determined by measuring optical density (OD_600_) on a Spark^®^Microplate reader (Tecan). The MIC was scored as the lowest antibiotic concentration where no growth was observed following subtraction of the OD_600_ of a well containing sterile medium.

### β-Galactosidase assays

For quantitatively assessing induction of *lacZ* reporter constructs in *E. faecalis*, exponentially growing cells (OD_600_ = 0.4-0.5) inoculated 1:250 from overnight cultures in BHI medium were exposed to different concentrations of bacitracin for 1 h or daptomycin for 30 minutes. Cells were harvested via centrifugation and stored at -20°C. β-Galactosidase activities were assayed in permeabilised cells and expressed in Miller units (MU) (Gebhard *et al*., 2006). For this, cells were resuspended in 1 ml Z-buffer (8.04 g Na_2_HPO_4_*7H_2_O, 2.76 g NaH_2_PO_4_*H_2_O, 0.123 g MgSO_4_*7H_2_O and 5 mL 1M KCl in 495 mL dH_2_O, pH 7). The samples were adjusted to OD_600_ = 0.5 in a 1 ml volume of Z-buffer and from this, two volumes were taken: 200 µl and 400 µl cells made up to 1 mL each with Z-buffer. This volume corresponds to the ‘volume of cells’ in the Miller Unit (MU) equation below. Following this, 20 µl 0.1% (w/v) SDS and 40 µl chloroform were added and vortexed for 5 seconds, then rested for 5-10 minutes. Reactions were started by adding 200 µl *o*-nitrophenyl-β-D-galactopyranoside (ONPG) (4 mg mL^-1^ in Z-buffer) and incubated at room temperature until yellow colouration was observed. If no colour change was visible, the reaction was incubated for 20 minutes. Reactions were stopped by adding 500 µl 1M Na_2_CO_3,_ and the time recorded, which corresponds to the ‘time’ in the Miller Unit (MU) equation below. Absorbance at 420 nm (A_420_) was then read. MUs were calculated using the following equation:

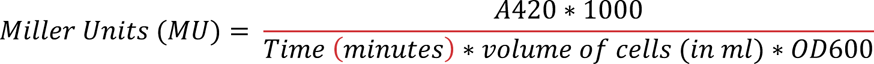

### Computational modelling

Rather than directly modelling the temporal dynamics of the regulatory network as done for the *B. subtilis* Bce system (Fritz *et al*., 2015; Piepenbreier *et al*., 2020), we chose to focus on the (meta) stable state reached in response to challenge with a given bacitracin concentration. The quantities modelled are the concentrations of bacitracin [*bac*], UPP [*UPP*], UPP-bound bacitracin [*UPPbac*], and the effective activities of SapRS [*SapRS*], RapAB [*RapAB*], LiaX [*Lia*], and DltXABCD [*Dlt*]. For the modelling, we made repeated use of a soft-switch sigmoid type function

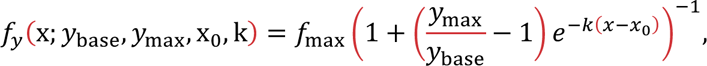

where x is the input quantity and y the output which varies between *y*_base_ and *y*_max_, with x_0_ controlling the switching threshold and k setting the sharpness of the transition. The equations of state for the model are then simply

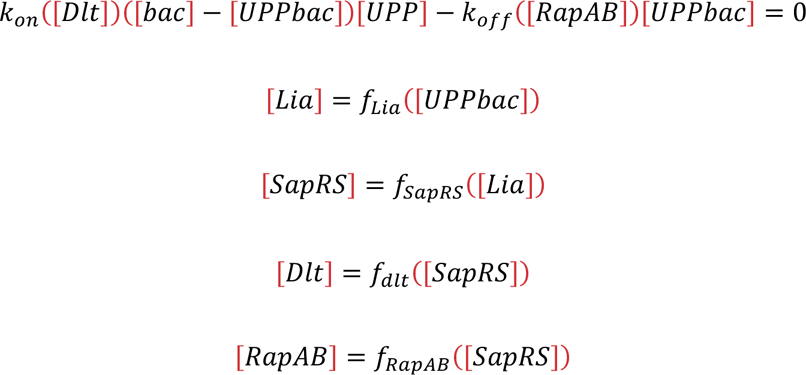

We modelled the binding and off rates for the UPP / bacitracin interaction as being linearly proportional to [*Dlt*] and [*RapAB*], respectively, with Dlt reducing bacitracin binding and RapAB increasing the off-rate. The biological reasoning for this is explained in the Results section.

For [*bac*], actual values from the experimental work were used, and the switching parameters for [*Dlt*] and [*RapAB*] as functions of [*SapRS*] were determined by fitting actual activity levels observed experimentally of these quantities for different bacitracin levels. The remaining parameters were fitted to achieve a description of experimental results for the wild-type strain by the model output. The values for each parameter are given in Table S3. Solving the equations of state for a given input bacitracin level [*bac*] yielded predictions for [*UPP*] and [*UPPbac*], which in turn drive the response curves plotted in Figure 3B.

### Data accessibility

The numerical data underpinning the results shown in Figures 1-4 are available in Table S4. The *Mathematica* file for the model is provided in the supplementary material.

## Acknowledgements

This work was supported in part by grant MR/N0137941/1 for the GW4 BIOMED MRC DTP, awarded to the Universities of Bath, Bristol, Cardiff and Exeter from the Medical Research Council (MRC)/UKRI. The authors gratefully acknowledge the Technical Staff within the Life Sciences Department at the University of Bath for technical support and assistance in this work. We would also like to thank Mathew Upton for the gift of *Streptococcus gallolyticus*.

## Author contributions

SMM, SG, TR and GF contributed to the conception of the study; SMM, LW and SG designed the experimental work; SMM, LW and OR carried out all data acquisition; SMM, LW, TR and SG carried out data analysis and interpretation; SMM, LW, SG, TR and GF wrote the manuscript.

## Graphical abstract

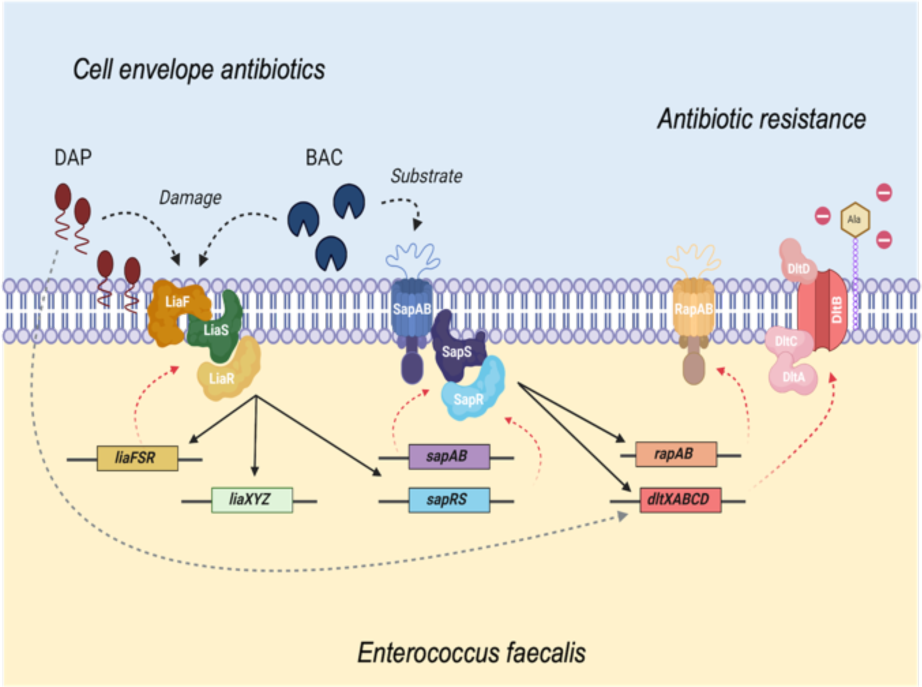

## Abbreviated summary

This study explored a regulatory network controlling resistance against daptomycin and bacitracin in *Enterococcus faecalis*. We show that resistance gene expression for both antibiotics is controlled by two regulatory systems, LiaFSR and SapRS, working in sequence and together processing two stimuli. Convergence of this network on the *dltXABCD* target operon, known to be associated with daptomycin resistance may explain why clinical resistance can emerge via regulatory mutations.

## Supplementary Material

**Table S1.** Vectors, plasmids and strains used in this study.

**Table S2.** Primers used in this study.

**Table S3.** Parameter values used in the mathematical model.

**Table S4.** Experimental numerical data for Figures 1-4.

**File S1.** Mathematical model file for the application *Mathematica*

**File S2.** Description, methods and results of interaction studies between antibiotic producing strains and *E. faecalis* wild type and deletion strains.

## File S2: Description, methods and results of interaction studies between antibiotic producing strains and *E. faecalis* wild type and deletion strains

### Results

#### The network components vary in importance during interactions with antimicrobial producer strains

As stated in the main manuscript, we were surprised by the complexity of the regulatory pathway controlling what in other Firmicutes bacteria is a fairly straightforward response to antibiotic challenge, where each regulatory system controls its own resistance genes. To shed some light on the reasons for the complexity of signalling in enterococci, we considered the environments these bacteria can be found in. *E. faecalis* is a common member of many natural microbial communities, such as in soil and water or the gastrointestinal tract of humans and animals. There, the enterococci reside within the small and large intestine and represent up to 1% of the faecal flora (Lorian, 1994; Sghir *et al*., 2000; Tendolkar *et al*., 2003; Eckburg *et al*., 2005). In such environments, *E. faecalis* interacts with other microbes and must defend itself against antimicrobial producing bacteria. Therefore, we wanted to assess the role of the individual components of the resistance network in protecting *E. faecalis* from antimicrobial activity produced by potential competitor bacteria. To do this, we used deferred antagonism assays to simulate relevant environmental pressures from other microbes the enterococci may encounter. We used four antimicrobial producing strains: the subtilin producer *Bacillus subtilis* ATCC6633 (Banerjee and Hansen, 1988), bacitracin producer *Bacillus lichenformis* ATCC10716 (Johnson *et al*., 1945), nisin-A producer *Lactococcus lactis* NZ9000 (Kuipers *et al*., 1998) and nisin-P producer *Streptococcus gallolyticus* AB39 (Aldarhami *et al*., 2020). Each producer was spotted onto a plate, and the antimicrobial was allowed to accumulate over multiple days. This was followed by the addition of an overlay containing the different *E. faecalis* strains to assess their sensitivity against the produced compounds based on the size of the zone of inhibition (Figure S1). Please note that we cannot confirm that each of these strains produced their expected antimicrobial under the chosen growth conditions. However, each producer caused the appearance of an inhibition zone in the *E. faecalis* overlay, showing that some antimicrobial compounds were indeed secreted.

Firstly, we tested the *E. faecalis* strains against the antimicrobial subtilin, produced by *B. subtilis* ATCC6633. Of the components under investigation in this study, the LiaFSR system appeared to contribute most strongly to resistance against subtilin, as the Δ*liaR* mutant displayed the largest increase in zone of inhibition compared to the wild type. The deletions of *sapR*, *sapAB* and *rapAB* also displayed an increased zone of inhibition compared to the wild type, but to a lesser extent than Δ*liaR*. This therefore suggests they play a lesser role in subtilin resistance.

In contrast, when testing the *E. faecalis* strains against the bacitracin producer *B. lichenformis* ATCC10716, deletion of *sapR, sapAB* and *rapAB* showed similar increased sensitivity, consistent with their contribution to the bacitracin resistance network that we have examined in this study and previously (Gebhard *et al*., 2014). Although also presenting increased sensitivity compared with the wild type, deletion of *liaR* resulted in a marginally smaller zone of inhibition compared with the other deletion strains. This is in line with our data presented above, showing the Lia system has a more indirect role in controlling the bacitracin response by regulating *sapRS* expression.

We next tested the *E. faecalis* mutants against the nisin producers, *L. lactis* NZ9000 and *S. gallolyticus* AB39. Although there was some inhibition of wild type *E. faecalis* by the nisin-A producer *L. lactis*, there was no difference in sensitivity between the wild type and deletion strains. This suggests either that the genes of our regulatory pathway do not contribute to resistance against this antibiotic, or that under the chosen conditions the inhibitory activity of *L. lactis* NZ9000 is primarily due to a different antimicrobial than nisin-A. In contrast, the nisin-P producer *S. gallolyticus AB39*, despite having no inhibitory effect on wild type *E. faecalis*, was able to strongly inhibit growth of the *liaR* deletion strain. Deletions of both *sapR* and *sapAB* also resulted in increased sensitivity but to a lesser degree than *liaR* deletion. Deletion of *rapAB* had little to no effect compared to the wild type, suggesting that the other target genes of the regulatory pathway but not the RapAB transporter are responsible for nisin P resistance.

These findings indicate that in response to different antimicrobials, the various members of this resistance network have differing roles to play and also differ in their relative importance. In response to subtilin and nisin-P, LiaR clearly plays a very important role in resistance, but it is a less important component in response to bacitracin exposure. This reflects LiaR’s role as a key regulator in the cell envelope stress response more widely, but with mostly a moderating role in response to bacitracin.

The *sapR*, *sapAB* and *rapAB* deletions all present with very similar effects in response to the antimicrobials, reflecting the interdependent functional relationship between the three components. The notable exception to this is RapAB in the context of nisin-P, where the transporter does not appear to contribute to protection of the cell.

**Figure S1.**
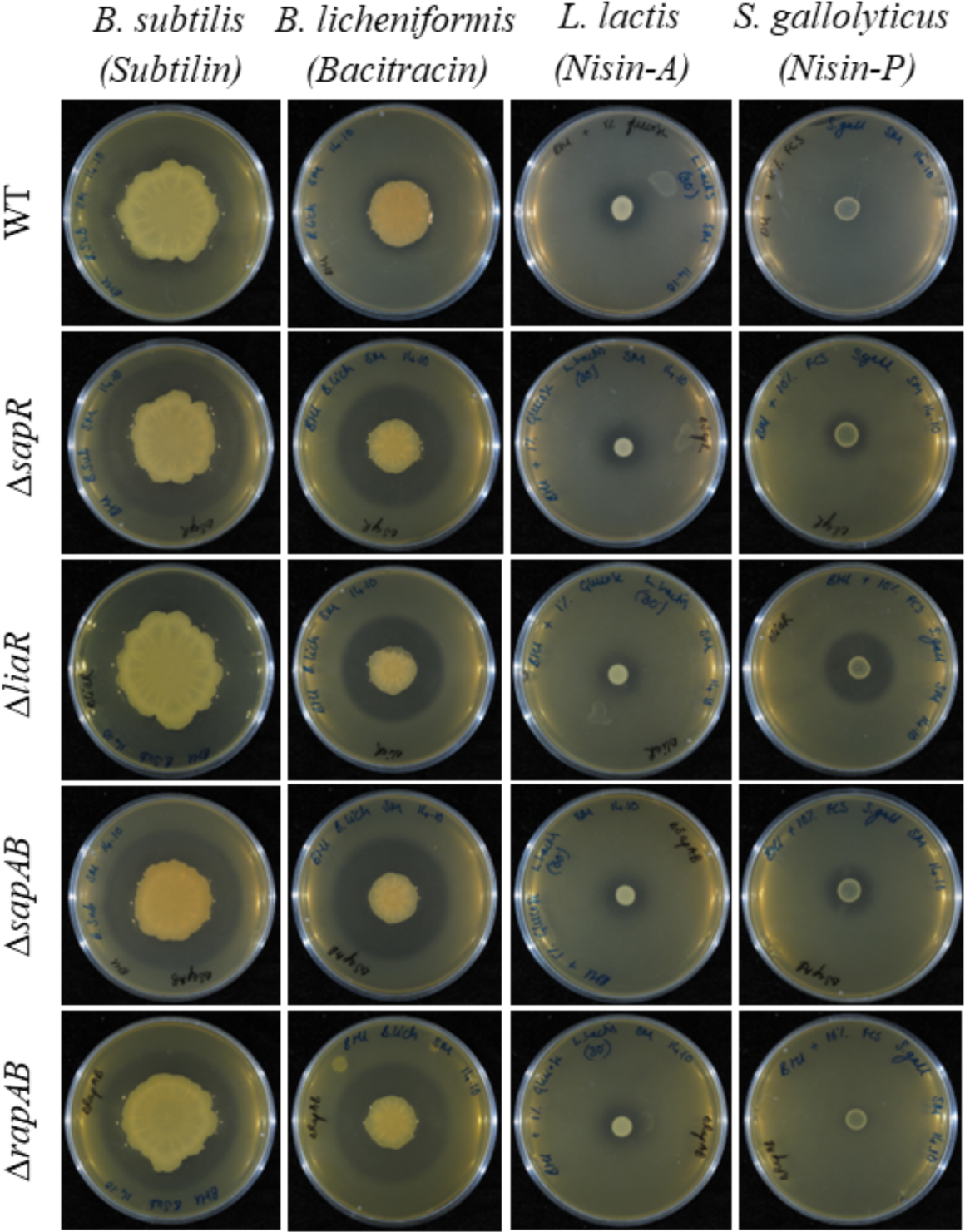
*E. faecalis* deletion strains show differential zones of inhibition in deferred antagonism tests against antimicrobials produced by other firmicute bacteria. The antimicrobial producer strains indicated at the top were grown overnight and then adjusted to an OD600 of 0.5 with fresh media. Aliquots (5ul) from each producer strain was then spotted and incubated at 25°C for 36-45 hours to allow antimicrobial accumulation. Overnight cultures of the *E. faecalis* strain indicated on the left were then added as a soft agar overlay. Plates were incubated at 25°C for 24 hours and zones of inhibition in the *E. faecalis* lawn were used to assess susceptibility to antimicrobials produced by the central strain.

### Experimental Procedures

#### Antagonism assays on solid media

A plate-based assay was utilised to measure growth inhibition between antibiotic producer strains and *E. faecalis* strains. For each species, cultures of the antimicrobial-producing bacteria were grown overnight in the respective growth medium and temperatures stated above and adjusted to OD_600_ = 0.5. Aliquots (5 µl) from each producer culture were spotted onto the centre of a BHI agar plate and incubated at room temperature (20-25°C) for 3-5 days to allow accumulation of antimicrobial products. Overnight cultures of *E. faecalis* strains were grown at 37°C in BHI medium, inoculated 1:100 into fresh medium and grown to OD = 0.5. Next, 3 mL of liquid BHI soft agar (7.5 g/L, 50°C) were inoculated with 30 μL culture. The soft agar was then poured onto the plate containing the pre-grown antimicrobial-producing strains and left to dry. The plate was then incubated overnight at room temperature to allow *E. faecalis* to grow to visualise the zone of inhibition. Results were recorded photographically using a PowerShot G camera attached to a lightbox.

---

**Table S1.**
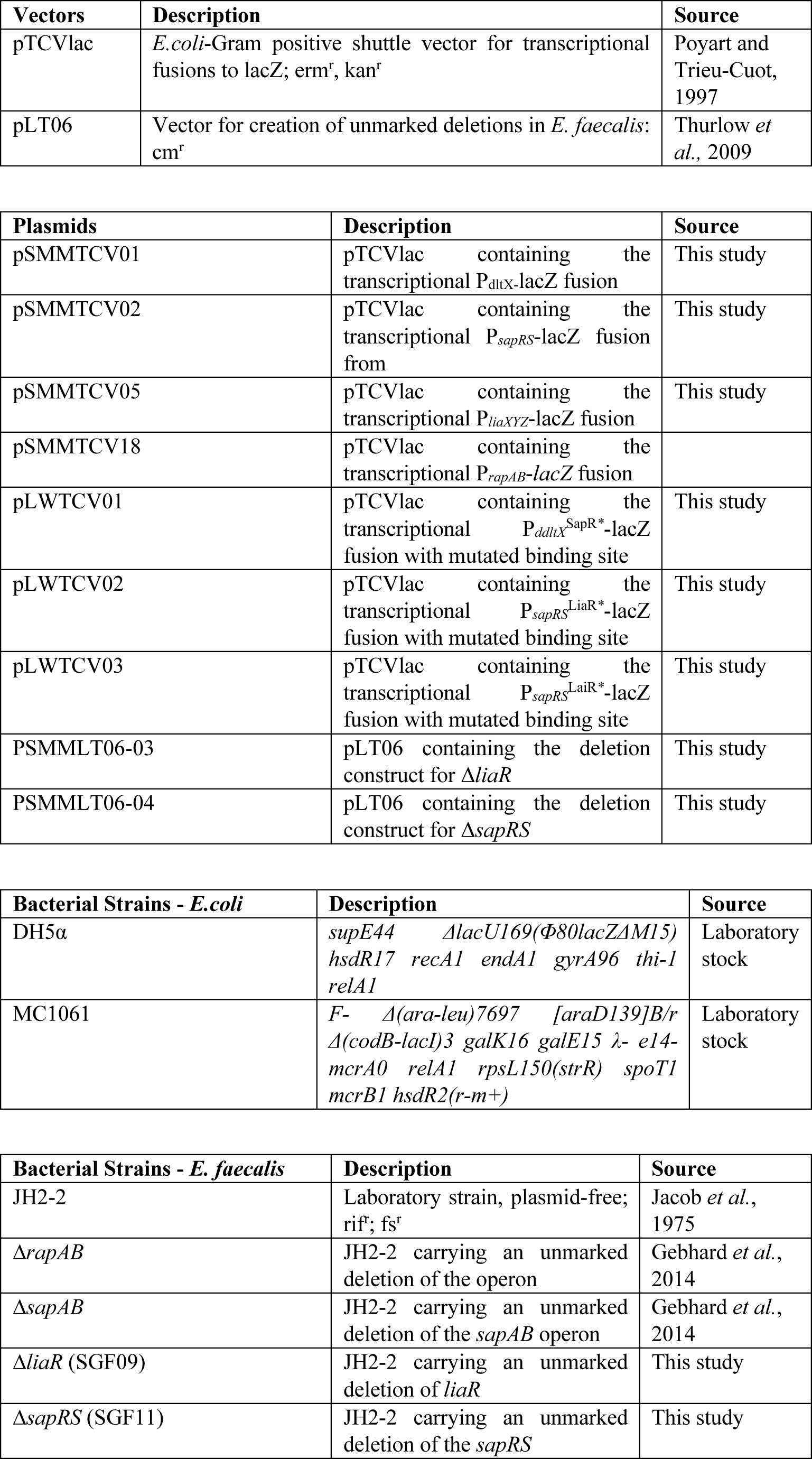
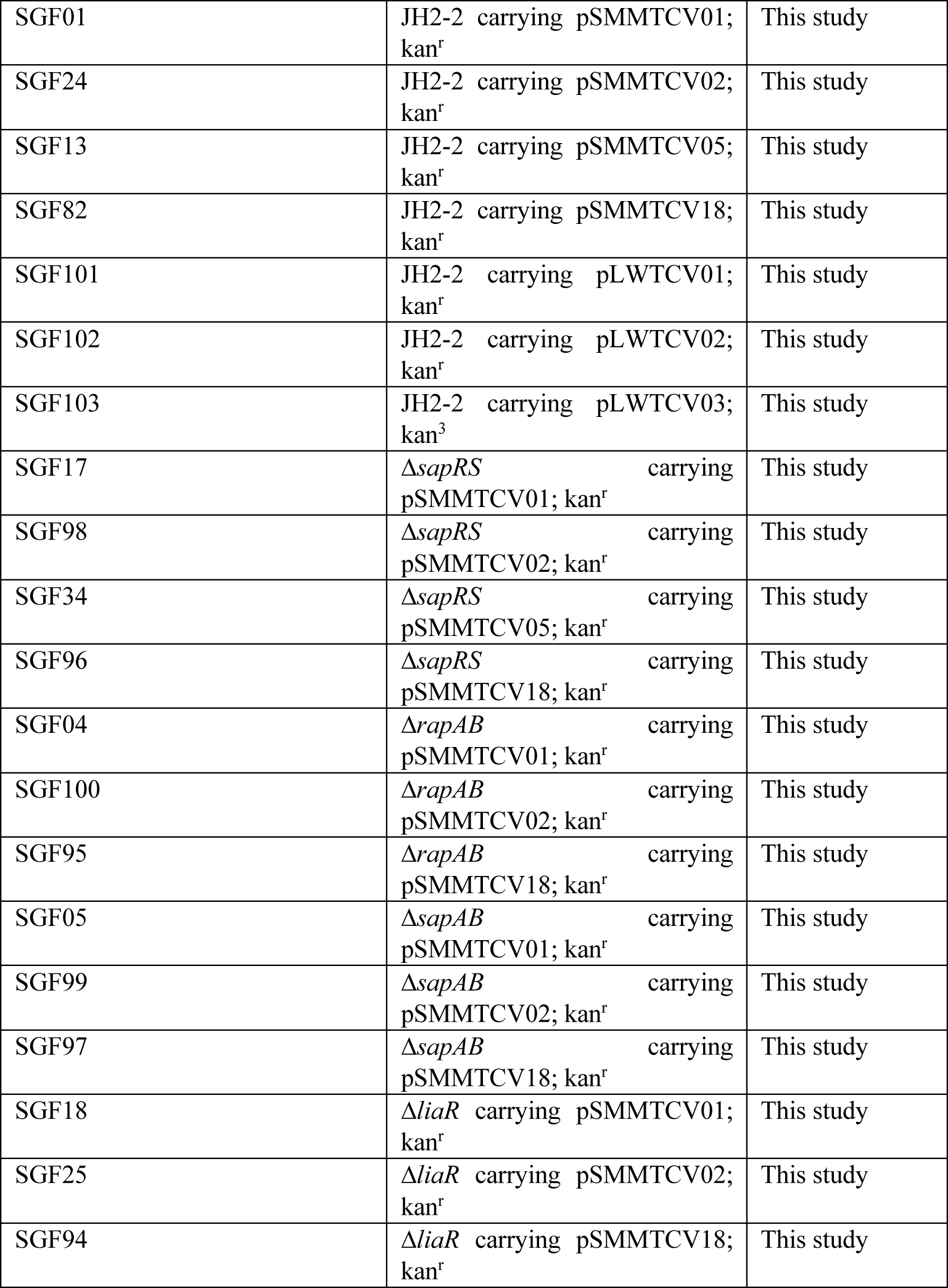
Vectors, plasmids and strains used in this study.

**Table S2.**
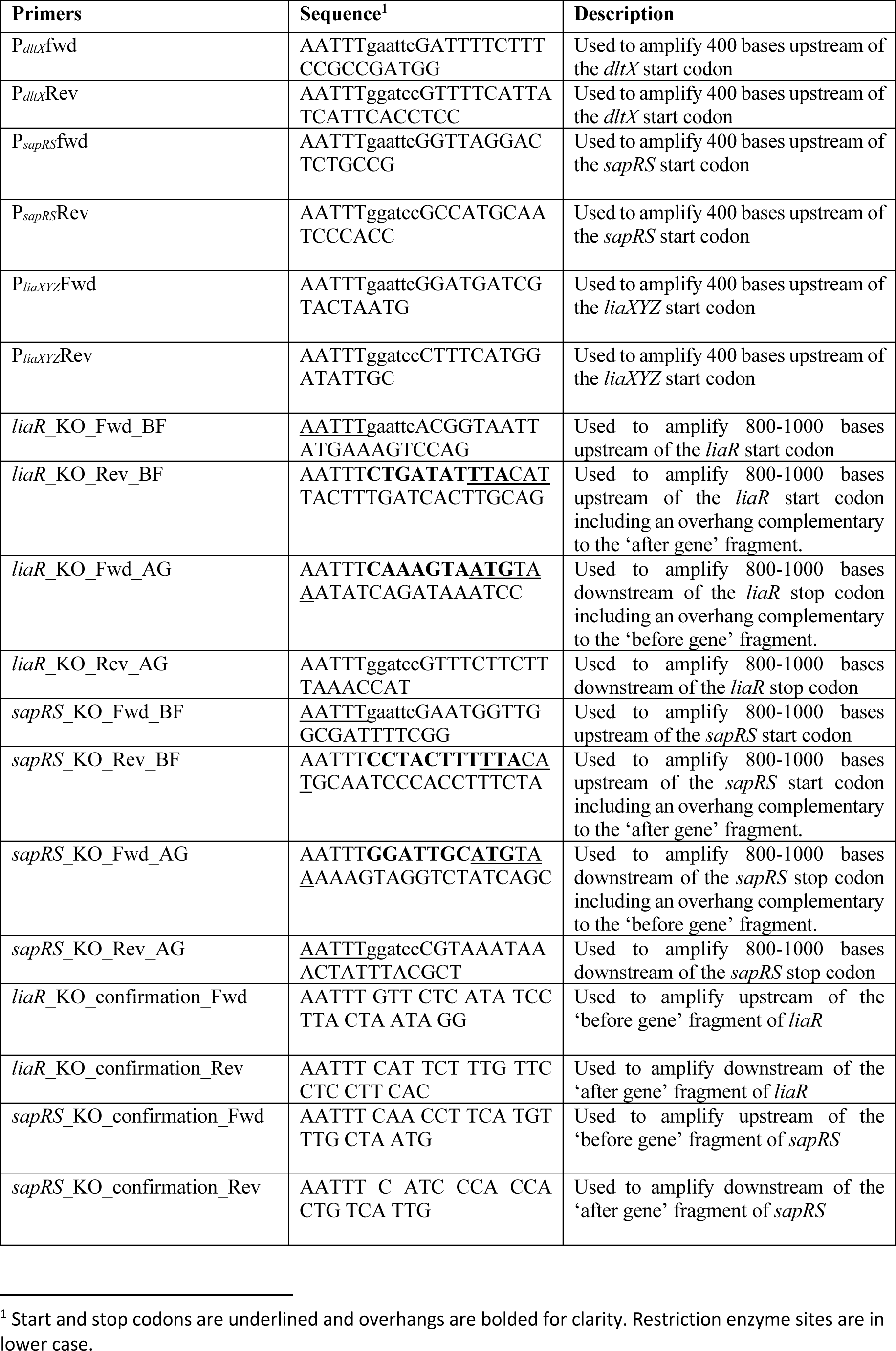
Primers used in cloning.

## Notes

<u>Funding statement:</u> This work was supported in part by grant MR/N0137941/1 for the GW4 BIOMED MRC DTP, awarded to the Universities of Bath, Bristol, Cardiff and Exeter from the Medical Research Council (MRC)/UKRI.

<u>Conflicts of interest:</u> The authors declare no conflicts of interest.

### Competing Interest Statement

The authors have declared no competing interest.

### Summary of Updates

New data are included in figures 1, 3 and 4. The corresponding text was re-written and/or amended and the discussion updated. Some material was moved to the supplemental section included at the end of the uploaded file. The title was changed and some minor text edits done.

